# Functional and structural modifications of influenza antibodies during pregnancy

**DOI:** 10.1101/2021.05.18.444722

**Authors:** Madeleine F. Jennewein, Martina Kosikova, Francesca J. Noelette, Peter Radvak, Carolyn M. Boudreau, James D. Campbell, Wilbur H. Chen, Hang Xie, Galit Alter, Marcela F. Pasetti

**Affiliations:** The Ragon Institute of MGH, MIT, and Harvard, Cambridge, MA 02139; Laboratory of Pediatric and Respiratory Viral Diseases, Division of Viral Products, Office of Vaccines Research and Review, Center for Biologics Evaluation and Research, US Food and Drug Administration, Silver Spring, MD, 20993; Center for Vaccine Development and Global Health, University of Maryland School of Medicine, Baltimore, MD 21201

**Keywords:** Pregnancy, antibodies, IgG, Fc-functionality, Fc-receptors

## Abstract

Pregnancy represents a unique tolerogenic immune state which may alter susceptibility to infection and vaccine-response. Here we characterized humoral immunity to seasonal influenza vaccine strains in pregnant and non-pregnant women. Pregnant women had reduced hemagglutinin subtype-1 (H1)-IgG, IgG1, and IgG2, hemagglutination inhibition and group 1 and 2 stem IgG. However, H1-specific avidity and FcγR1 binding increased. Influenza-antibodies in pregnancy had distinct Fc and Fab glycans characterized by di-galactosylation and di-sialylation. In contrast, agalactosylation and bisection were prominent outside of pregnancy. H1-specific Fc-functionality was moderately reduced in pregnancy, although likely compensated by stronger binding to cognate antigen and FcR. Multivariate analysis revealed distinct populations characterized by FcγR1 binding, H1-IgG levels, and glycosylation. Pooled sera from pregnant women exhibited longer retention *in vivo*. Our results demonstrate structural and functional modulation of humoral immunity during pregnancy in an antigen-specific manner towards reduced inflammation, increased retention in circulation, and efficient placental transport.

## INTRODUCTION

The immune system adapts in a precise and unique way during pregnancy to support fetal development and facilitate full-term delivery. These adaptations involve changes in cell phenotypes and frequencies (Th2 polarization, reduction of circulating NK and T cells, reduction in the number of B cells) as well as modulation of cell function (i.e., cytokine production) (Kraus et al. 2010; Kraus et al.2012). In addition, antibodies produced during pregnancy display unique glycosylation patterns (Bondt et al. 2014; Bondt et al. 2013; Einarsdottir et al. 2013). Collectively, these changes favor a non-inflammatory and tolerogenic environment. As a result, auto-immune conditions, such as rheumatoid arthritis, may transiently resolve during pregnancy (Kraus et al. 2010; Memoli et al. 2013). It has been proposed that a corollary of these alterations is increased maternal susceptibility to pathogens, which would also affect fetal health and development. For example, pregnant women are known to experience more severe influenza virus infection and are at higher risk of death, as evidenced during the 2009 H1N1 pandemic (Kourtis, Read, and Jamieson 2014). During the COVID-19 pandemic, symptomatic maternal infection posed a higher risk for unfavorable outcomes requiring hospitalization and invasive ventilation (Zambrano et al. 2020) and for fetal complications, including preterm birth, growth restriction, and miscarriage (Yee et al. 2020; Barrero-Castillero et al. 2020). These can be mitigated by interventions that strengthen prenatal immunity.

There is a long history of safe and effective immunization of pregnant women against tetanus and diphtheria (Forsyth et al. 2015). In the past decade, the introduction of maternal acellular pertussis vaccine (ACIP 2013) propitiously reduced infant hospitalizations and respiratory infections (Ohfuji et al. 2018; Regan et al. 2016). Seasonal influenza vaccination during pregnancy has been recommended to reduce the risk of infection and severity of disease of mothers and young infants (ACIP 2013; Fiore et al. 2010; Safety 2014). Several studies have demonstrated efficacy of this approach in reducing both maternal and infant hospitalization and illness (Thompson et al. 2019).

Recognizing the value of this strategy to improve public health, a growing number of vaccine candidates intended to strengthen maternal-infant immunity through vaccination during pregnancy are advancing in the clinical pathway (Saso and Kampmann 2020; Vojtek et al. 2018).

Conceivably, the profound physiological changes required to sustain pregnancy may also influence vaccine-induced immunity. Clinical studies of vaccination in pregnancy have focused primarily on the safety of these new interventions and their immunogenicity, through the analysis of serum antibody levels in mothers and their infants (Monath and Nasidi 1993; Sperling et al. 2012; Schlaudecker et al. 2012; Ohfuji et al. 2018; Munoz et al. 2020). A few of these studies have examined maternal T cell responses (Huygen et al. 2015) and/or functional capacity of vaccine-induced antibodies (Jennewein et al. 2019). However, it remains largely undefined whether the state of pregnancy *per se* influences vaccine-induced adaptive immunity and if that is the case, what the implications of such changes would be for both maternal and infant health.

Considering that antibodies are essential for protection against influenza, the goal of this study was to examine the biophysical and functional properties of influenza-specific antibodies in pregnant and non-pregnant women who received seasonal influenza vaccination. We tested the hypothesis that pregnancy-associated structural changes would likely affect both Fab- and non-neutralizing Fc-antibody mediated functions, which are key contributors of protective immunity (Jennewein et al. 2019). To this end, we conducted an in-depth characterization of antibodies to matched seasonal influenza vaccine strains in pregnant and non-pregnant women that included: 1) serum antibody levels determined by hemagglutinin (HA) globular head and stem-specific IgG ELISA, hemagglutination inhibition (HAI) and microneutralization (MN); 2) HA-specific IgG avidity; 3) HA-specific IgG subclasses and Fc receptor binding by ELISA; 4) glycan profile of total and HA-specific IgG Fc and Fab molecules; 5) Fc-mediated innate cell functions: antibody dependent neutrophil phagocytosis (ADNP), antibody dependent monocyte phagocytosis (ADCP), antibody dependent complement deposition (ADCD), and antibody dependent NK cell degranulation; and 6) *in vivo* analysis of maternal IgG retention. Immunological outcomes in the two groups were compared using multivariate analysis. Distinct profiles of influenza-specific IgG Fab and Fc glycan profiles and function were identified between pregnant and non-pregnant vaccine recipients, with subtype-specific antibodies being differentially affected. Two well-defined segregated clusters were identified by aggregate data analysis. These results support similar studies to inform vaccine development and implementation in this important population.

## RESULTS

### Pregnancy associated differences in influenza-specific antibody titer

To understand pregnancy-associated modulation of humoral immunity, we conducted a comparative analysis of the magnitude, structural features, and functionality of influenza antibodies in pregnant and non-pregnant women after seasonal influenza vaccination **(Supplemental Table 1)**.

The breadth of antibody responses was determined through analysis of full-length HA- and HA-stem-specific serum IgG titers **(Figure 1A, B)** and canonical function of influenza antibodies: hemagglutinin inhibition (HAI) and microneutralization (MN) **(Figure 1C, D)**. Antibody reactivity was tested against two of the subtypes included in the vaccine 2017-18 northern hemisphere flu vaccine ((WHO) 2017): H1 MI (drifted strain of the 2009 H1N1 pandemic lineage which was included since the 2016-17 Northern Hemisphere vaccine--A/Michigan/45/2015, H1N1) and H3 HK (A/Hong Kong/4801/2014, H3N2), as well as against the original 2009 H1N1 pandemic strain H1 CA (A/California/07/2009, H1N1) that disproportionally and severely affected pregnant women during the 2009 pandemic (Jamieson et al. 2009; Louie et al. 2010).

**Figure 1:**
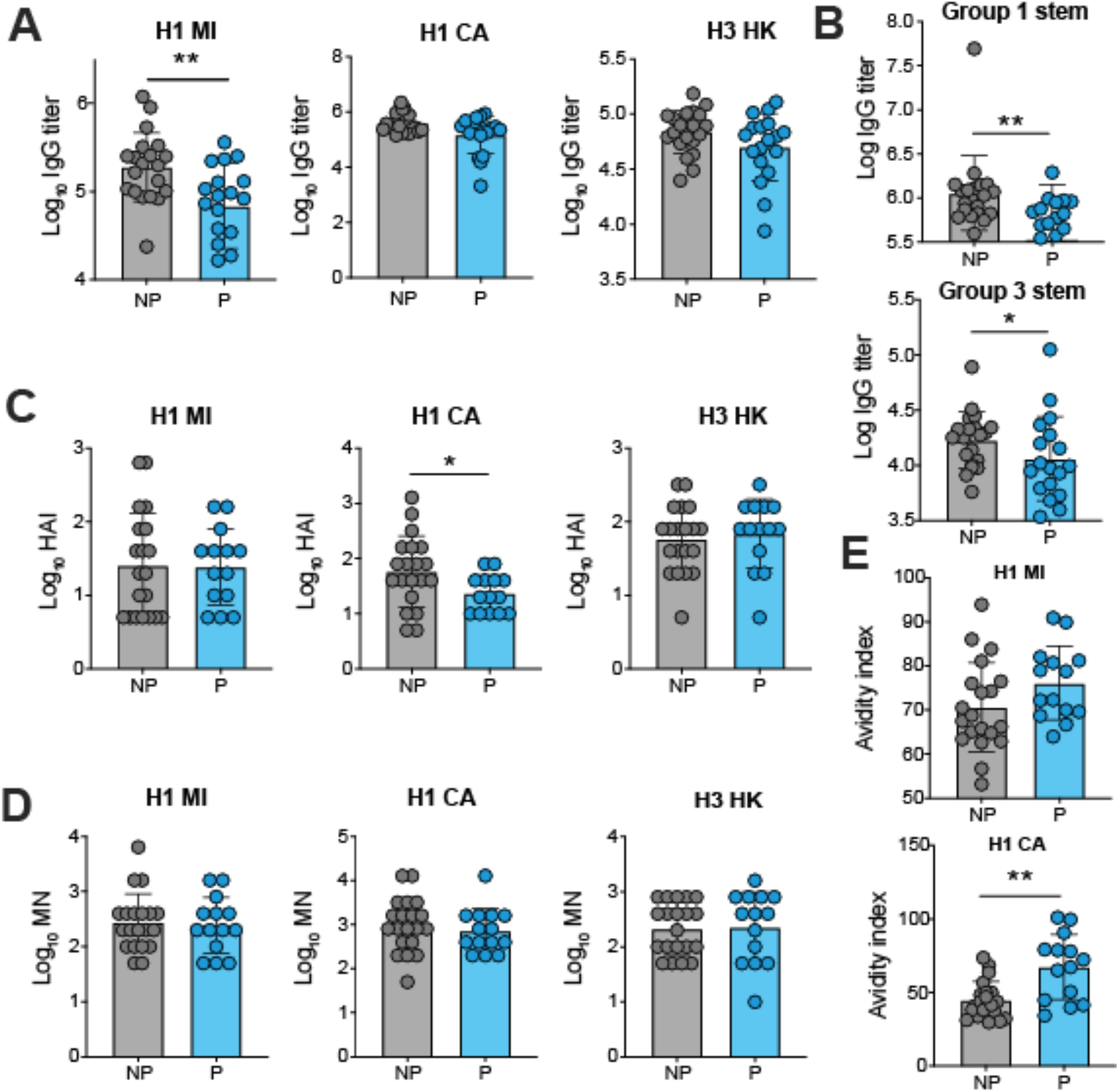
Pregnancy associated differences in influenza-specific antibody titer. HA-specific IgG **(A)**, group 1- and group 3-stem specific IgG **(B)**, hemagglutination inhibition (HAI) **(C)**, microneutralization (MN) **(D)** and HA-specific IgG avidity **(E)** in pregnant and non-pregnant women. Data represent log transformed individual datapoints and geometric mean titers (bars). ELISA, HAI and MN titers were compared using one-tailed Mann-Whitney test. Avidity was evaluated using a two-tailed Mann-Whitney test *p<0.05, **p<0.01.

Pregnant women had significantly lower IgG titers against H1 MI HA, as compared to the non-pregnant, while titers against H1 CA were unaffected **(Figure 1A)**. A trend of lower H3-IgG was also observed **(Figure 1A)**. Serum IgG directed to the conserved group 1 and group 2 stem regions were both substantially reduced in the pregnant women **(Figure 1B)**. Although both groups had similar HAI and MN titers for the contemporary vaccine strains H1 MI and H3 HK **(Figure 1C, D)**, pregnant women did exhibit significantly reduced HAI titers against the original pandemic strain H1 CA **(Figure 1C)**.

Intriguingly, the avidity of IgG against H1 CA HA was higher in the pregnant women as compared to the non-pregnant **(Figure 1E)**. Likewise, a trend of higher IgG avidity against H1 MI HA was observed. Together, these results hint at pregnancy-associated differences in magnitude, canonical functions, and strength of binding of influenza antibodies, which appear to be antigen-specific.

### Pregnancy influences IgG subclass and Fc receptor binding

To better understand the profile of influenza-specific antibodies elicited during pregnancy, we conducted a detailed class (IgG, IgA, and IgM) and subclass (IgG 1-4 and IgA (1-2) analysis of serum antibodies specific for full-length Has H1 MI and H3 HK and their binding capacity to different Fc receptors using a multiplex luminex assay **(Figure 2)**. Consistent with the ELISA data described above **(Figure 1A)**, pregnant women had significantly reduced levels of H1 MI HA-specific total IgG, IgG1, IgG2, and IgG3 as compared to non-pregnant women **(Figure 2A, B)**. A similar trend of lower antibody levels (as seen previously by ELISA, in Figure 2) remained for H3 HA-specific IgG and IgG1, and no differences were seen in either H1 MI or H3 HA-specific IgA1, IgA2, or IgM levels between the two groups **(Figure 2A, B)**. Pregnant women had a significantly lower ratio of IgG1 and IgG3 to IgG4 for both antigens **(Figure 2C, D)**.

**Figure 2:**
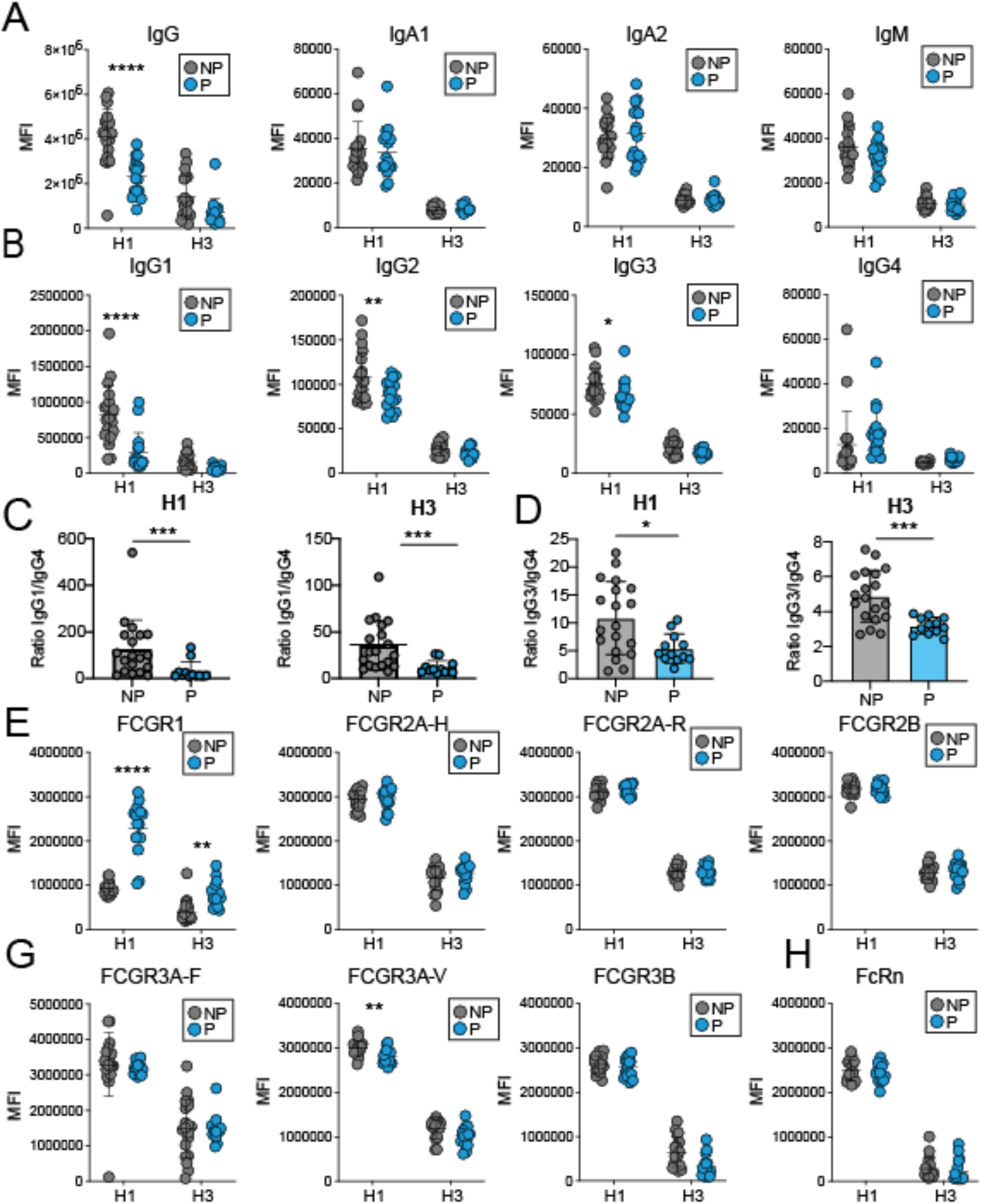
Pregnancy influences HA-specific IgG subclass and Fc receptor binding. IgG and IgG subclass levels and the Fc receptor binding were assessed by Luminex. **A-B**. Dot plots depict the differences between isotype and subclass levels for non-pregnant (NP, grey circles) and pregnant (P, blue circles) women. **A**. Isotypes. **B**. IgG subclasses. **C-D**. IgG1/IgG4 **(C)** IgG3/IgG4 ratios (D). **E-H**. Class 1 Fc receptor. **D**. Class 2 Fc receptors. **E**. Class 3 Fc receptors. **F**. Neonatal Fc receptor. Statistics evaluated as Mann-Whitney test and stars represent differences between H1 and H3 response.*p<0.05, **p<0.01, ***p<0.001, ****p<0.0001.

Clear differences were also seen in terms of HA antibody binding to Fc-receptors between the two groups **(Figure 2E-H)**. Unlike HA-specific antibody binding patterns revealed by ELISA or luminex assays, both H1 MI HA- and H3 HK HA-specific antibodies from pregnant women exhibited increased binding to FcγR1 as compared to the non-pregnant **(Figure 2E)**. No differences in antibody binding capacity to other FcγR molecules were detected between two groups **(Figure 2F-H)**, except for reduced H1 HA-specific IgG binding to FcγR3 allotypic variant V in the pregnant group **(Figure 2G)**.

### Pregnancy alters antibody Fc and Fab N-glycosylation

To identify pregnancy-associated changes in influenza-specific antibody glycosylation, we examined the Fc- and Fab-glycan composition of total and H1 MI- and H3 HK-specific serum IgG in both pregnant and non-pregnant women. For each of the samples, a total of 24 distinct glycan peaks were captured by capillary electrophoresis, and overall changes in the four differentially added sugars (galactose, sialic acid, bisecting n-acetyl glucosamine–GlcNAc–and fucose) were calculated for bulk **(Figure 3A, B)** and HA-specific **(Figure 3C-F, G-V)** IgG for each group.

**Figure 3:**
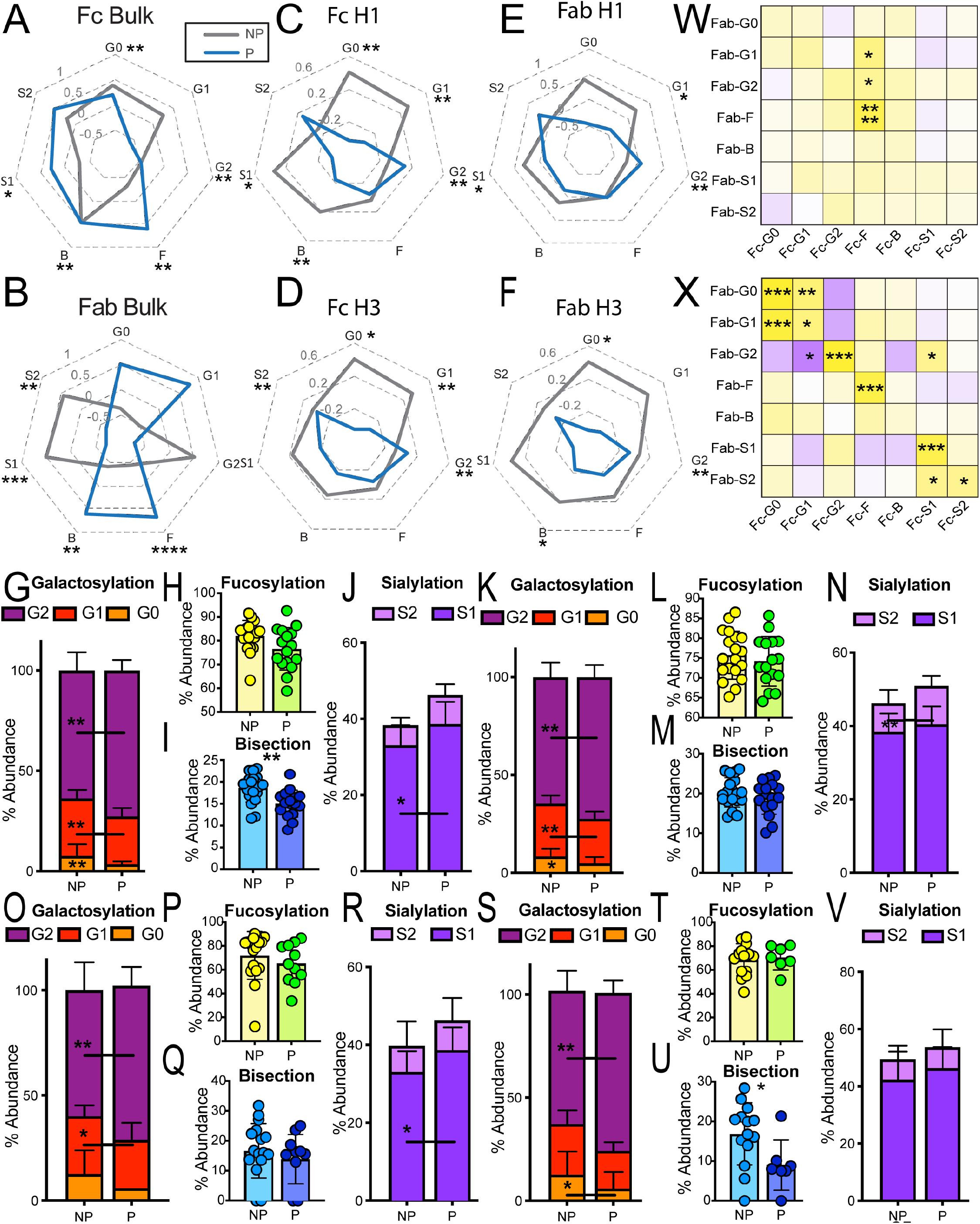
Pregnancy alters antibody Fc and Fab glycosylation. Antibody glycosylation for HA-specific and bulk antibodies was analyzed by capillary electrophoresis. **A-R**. Radar plots depicting different glycosylation profiles of bulk IgG Fc **(A)** and Fab **(B)**, H1-specific IgG Fc **(C)** and Fab **(E)** and H3-specific IgG Fc **(D)** and Fab **(E)** in non-pregnant and pregnant groups. G=galactose (0-2), F=fucose, B=Bisecting GlcNAc, S=sialic acid (1-2). **G-V**. These dot-line and stacked-bar plots depict the differences in HA1-specific antibody Fc **(G-J)** and Fab **(O-R)** and HA3-specific antibody Fc **(K-N)** and Fab **(S-V)** glycosylation between non-pregnant (NP) and pregnant (P) women. **(W, X)** Heatmaps showing the spearman correlation between HA-specific Fab and Fc glycosylation for pregnant **(W)** and non-pregnant **(X)** women. Significance for radar plots and glycosylation determined by Mann-Whitney test. *p<0.05, **p<0.01, ***p<0.001, ****p<0.0001.

Compared to bulk IgG of non-pregnant women, the bulk Fc glycosylation profile of pregnant women revealed shifts towards increased fucosylation (F) and sialylation (S1 and S2) consistent with an anti-inflammatory phenotype **(Figure 3A)**. On the other hand, pregnant women’s bulk Fab had fully fucosylated (F), agalactosylated (G0 and G1) and bisected (B) glycans **(Figure 3B, O-V)**.

In contrast to the bulk molecules, Fc and Fab glycans of HA-specific antibodies were largely concordant and included digalactosylated (G2) structures with increased sialylation and reduced bisection **(Figure 3C-F, G-V)** in the pregnant group, whereas agalactosylated, monosialylated and bisected structures were predominant in the non-pregnant group. Taken together, these results point to a distinct Fab and Fc glycan profile of antigen-specific antibodies produced during pregnancy consistent with distinct capacity for antigen-recognition (Fab) and finely modulated inflammatory properties (Fc).

To explore possible associations between Fab and Fc glycosylation, we performed correlation analyses between glycans of HA-specific antibodies in both groups **(Figure 3W, X)**. Significant associations between specific glycan types were detected **(Figure 3W, X)**, which suggest coordinated glycosylation of both Fab and Fc molecules. This was particularly evident in the non-pregnant group.

### Antigen-specific non-neutralizing antibody functions are selectively affected by pregnancy

Given the observed differences in magnitude and glycan composition of HA-specific antibodies during pregnancy, we next investigated their Fc mediated effector functions. Responses were measured independently for the two vaccine strains (H1 MI and H3 HK). Monocyte phagocytosis (ADCP) and complement deposition (ADCD) activity of H1 HA-specific antibodies were significantly lower in pregnant women as compared to the non-pregnant women **(Figure 4B, C)**. Intriguingly, no differences were seen in neutrophil phagocytosis (ADNP) and NK degranulation **(Figure 4A, D, E)**, Both H1 HA- and H3 HA-specific antibodies in the pregnant group exhibited reduced cytotoxicity (ADCC) although it did not reach statistical significance **(Figure 4F)**. Overall, a consistent trend towards reduced antibody functionality across FcR-dependent functions was observed in the pregnant women. The predominant changes on H1 HA-specific antibody function further supports pregnancy-associated impact on antigen-specific humoral immunity.

**Figure 4:**
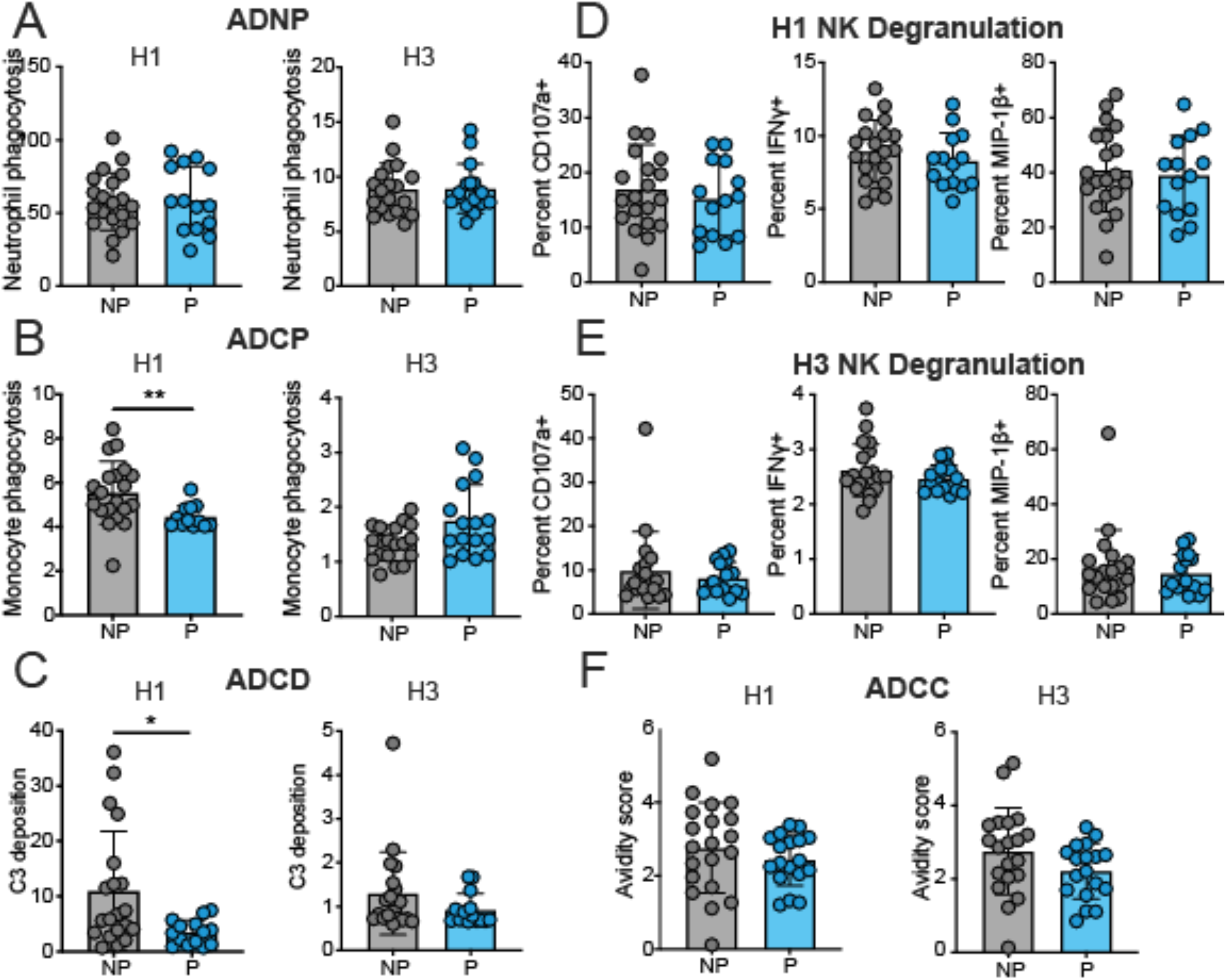
Antigen-specific non-neutralizing antibody functions are affected by pregnancy. Influenza H1- and H3-specific antibody dependent neutrophil phagocytosis (ADNP) **(A)**, monocyte phagocytosis (ADCP) **(B)**, complement deposition (ADCD) **(C)**, NK degranulation measured as CD107a, MIP-1β and IFNγ **(D, E)** and cellular cytotoxicity (ADCC) **(F)**. Groups were compared using two-tailed Mann-Whitney Test. *p<0.05, **p<0.01.

### Multivariate signatures of antibodies during pregnancy

To dissect the differences between antibodies in pregnant and non-pregnant women, a least absolute shrinkage and selection operator and LASSO Partial Least Squares Discriminant Analysis (LASSO-PLSDA) was conducted. This modeling technique allowed the determination of features that contributed most strongly to the group differences, and these predictors were then visualized on the PLSDA plot. This analysis included all the influenza-specific variables assessed (serum IgG, HAI and MN titers, glycosylation, Fc-effector functions, antibody subtypes Fc receptor binding), 79 in total. Readouts from pregnant and non-pregnant groups were separated and orthogonalized along LV1 **(Figure 5A)**. The down-selected features were ranked by their importance in separating the pregnant and non-pregnant antibody profile **(Figure 5B)**. The top predictor was H1 HA-specific antibody binding to FcγR1, which was greatly enriched in pregnant women. Another predictor was the lower H1 HA-specific IgG titer in the pregnant group. The remaining predictors were mainly Fc and Fab glycan residues. The groups segregated in antigen-specific antibody galactosylation profiles; anti-inflammatory H3 HA Fc G1B and H1 HA Fc G2S2F glycans were preeminent in pregnant women whereas inflammatory H1 HA Fc-G1F/G1FB and H1 HA Fab-G0 dominated in the non-pregnant group. Thus, Fc glycosylation and Fc receptor binding were the primary driving forces in determining the model. Most of the parameters down-selected with VIP scores were H1MI HA-specific even though H1 MI, H1 CA, and H3 HK HA-specific readouts were included in the model, reinforcing the notion of antigen dependency of pregnancy-induced antibody changes.

**Figure 5:**
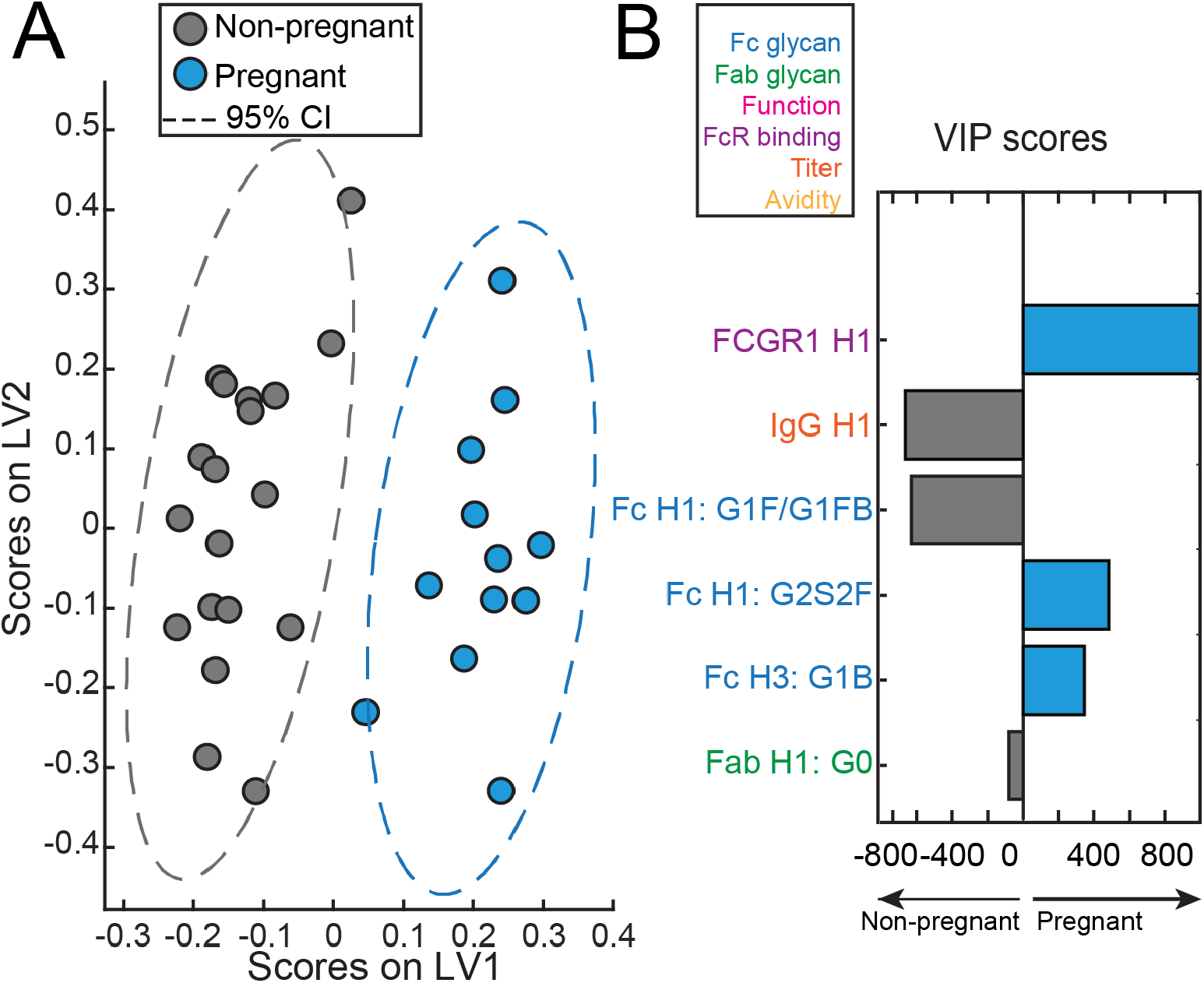
Multivariate signatures of antibodies produced during pregnancy. Computational analysis was used to distinguish features enriched in pregnancy. **A**. LASSO Partial Least Squares Discriminant Analysis (LASSO-PLSDA) of influenza-specific antibody features between pregnant and non-pregnant women orthogonalized along latent variable 1 (LV1). LV1 explains 42.5% of the variance along the X axis while LV2 explained 17.1% of the variance. **B**. The Variable Importance in Projection (VIP) scores for the PLSDA indicate the prime factors driving the differences between pregnant and non-pregnant women. Factors pointing towards the left are enriched in non-pregnant samples, while those pointing right are enriched in pregnant samples. Bar color and length corresponds to relative importance. Antibody features are colored by category.

### In vivo retention of antibodies produced during pregnancy

The distinct glycosylation of influenza antibodies in pregnant women begs the question of the purpose of such post-translational modifications, assuming it would ultimately confer some evolutionary advantage to support offspring’s health. Our group had shown selective transfer of maternal antibodies with digalactosylated Fc-glycan (Jennewein et al. 2019). These antibodies preferentially bind the FcRn receptor, which not only mediates translocation of IgG across the placenta but also recycling. Thus, we hypothesized that the concomitant increased galactosylation and sialylation of influenza-specific antibodies in pregnant women could allow longer antibody persistence. To address this question, we conducted an adoptive transfer experiment. Pooled serum from both pregnant and non-pregnant women were administered to mice, and antibody levels measured in circulation **(Figure 6)**. Despite starting at the same levels in the recipient mice, IgG from the pregnant group exhibited significantly higher retention in mouse circulation as compared to IgG from the non-pregnant group **(Figure 6)**. This is consistent with the hypothesized prolonged persistence and reduced catabolism.

**Figure 6.**
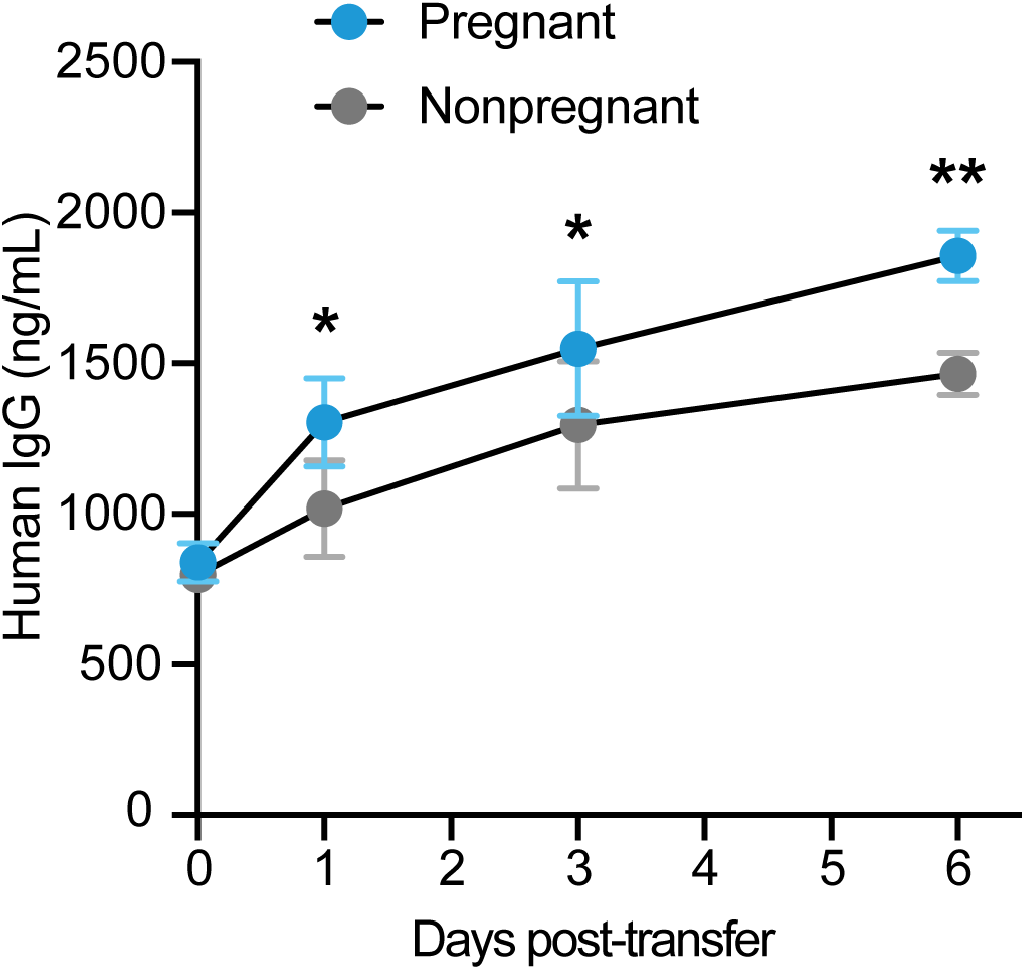
In vivo antibody retention. The dot-line plot depicts the retention of IgG in mice injected with pooled plasma from pregnant (blue, 4 mice) or non-pregnant (grey, 5 mice) mice. Retention was measured over a 6-day period. Dots are at mean with error bars at the standard deviation. Statistics evaluated using a two-way ANOVA. *p<0.05, **p<0.01.

## Discussion

Seasonal influenza vaccination during pregnancy is recommended by the Advisory Committee on Immunization Practices (Grohskopf et al. 2020) and the American College of Obstetricians and Gynecologists (“ACOG Committee Opinion No. 732: Influenza Vaccination During Pregnancy” 2018). The heightened susceptibility of pregnant women to severe influenza infection during the 2009 H1N1 pandemic (Siston et al. 2010) brought attention to the importance of vaccination for protection of both mothers and newborns. There has been renewed interest, in recent years, in expanding maternal immunization against other pathogens (e.g., respiratory syncytial virus, cytomegalovirus, group B streptococcus, etc.) that are important causes of neonatal and infant death. This approach, however, faces unique challenges. Fine-tuned regulatory mechanisms are deployed during pregnancy to balance the tolerogenic environment necessary to avoid unhealthy reactions to paternal antigens with the need to engage robust adaptive immunity against microbial threats (Mor and Cardenas 2010; Schumacher, Costa, and Zenclussen 2014). Profound changes in humoral immunity occur during gestation, which include decreased number of peripheral B cells (Watanabe et al. 1997; Faucette et al. 2015), and increased IL-10-secreting regulatory B cells (Rolle et al. 2013). These changes are also expected to impact vaccine-induced immunity.

In this study we report, for the first time, distinct binding and functional features of antibodies specific for seasonal influenza vaccine strains produced during pregnancy. Among the most notable is the overt reduction in total IgG and IgG subclasses specific for the H1 MI vaccine strain. HAI titers against H1 CA – the original 2009 H1N1 pandemic strain, were also reduced and, alarmingly, below seroprotective levels. Changes in antibody levels affected primarily those against H1 HA but not those against H3 HA. This could be due to immune imprinting, since H1 CA-like viruses have continued to circulate globally since the 2009 H1N1 pandemic (H1 MI is an antigenically drifted H1 CA-like virus) while the subclade of predominant H3 virus has changed almost every other year (WHO ; Allen and Ross 2018). The history of prior influenza exposure of these women and pre-existing immunity are not known and could have influenced the responses detected (Kosikova et al. 2018). Others have reported reduced HA-specific IgG and IgG subclass responses, as well as HAI titers to seasonal influenza vaccines during pregnancy (Schlaudecker et al. 2018; Schlaudecker et al. 2012; Bischoff et al. 2013). These studies had access to pre-vaccination samples and reported no differences in basal antibody titers between pregnant and non-pregnant women, and yet were able to detect similar reductions in post vaccine immunity, suggesting that the antibody modulations we have observed are intrinsic of pregnancy.

Pregnant women had significantly lower IgG1/IgG and IgG3/IgG4 ratios in response to both H1 and H3 vaccine strains, which is consistent with a Th2-type shift of humoral immunity during gestation. IgG1 and IgG3 are the main subclasses produced in response to viral antigens (Hjelholt et al. 2013), hence their reduced proportions raise concerns about insufficient protection against viral infections.

In addition to lower IgG response to the H1 vaccine strain, pregnant women had reduced IgG titers against highly conserved group 1 and group 2 HA stem regions. This observation is new and suggests that production of stem-specific antibodies during pregnancy might be impaired and that these antibodies could be limited in their capacity to recognize emerging virus variants. Further studies to understand HA stem-specific immunity in this group are warranted as stem antibodies are hypothesized to confer broad cross-protection (Wu and Wilson 2017) and are candidate antigens for sought after universal flu vaccines (Sautto, Kirchenbaum, and Ross 2018).

Intriguingly, we found that H1 HA-specific IgG in pregnant women had higher avidity compared to those from non-pregnant women. In addition, HA-specific antibodies produced during pregnancy had increased capacity to bind FcγR1. FcγR1 is an activating receptor expressed in DC and monocyte/macrophages, with high affinity for human monomeric IgG, particularly IgG1 (Stewart et al. 2014; Bruhns et al. 2009). On the other hand, H1 HA-specific IgG from pregnant women had reduced binding capacity to FcγRIIIa-V, a high affinity allotype expressed in monocytes/macrophages. The enhanced capacity of antibodies to bind cognate antigens and to selectively engage in innate cell function via Fc-interaction may represent compensatory mechanisms to maximize response yet avoid harmful effects.

Another novel aspect of our study was the analysis of extra-neutralizing FcR-mediated antibody functions in pregnant women. H1-specific ADCP, ADCD, and ADCC were reduced in this group, and this observation is consistent with reduced levels of H1 IgG, and in particular IgG1, which triggers these processes efficiently by engaging FcγRI (Gunn and Alter 2016). The reduced capacity of H1 HA antibodies to bind to FcγRIII-V, which is expressed in monocytes and macrophages might have also contributed to this outcome. IgG1 has been implicated in activation of complement in the context of influenza infection, hence reduced IgG1 in this group is consistent with lower C3 deposition.

Unlike FcR-mediated monocyte/macrophage uptake, neutrophil phagocytic activity and NK degranulation both appeared largely unaffected during pregnancy. Neutrophil phagocytosis can be activated by different and additional signals such as IgG2 interaction with FcγRIIa and IgA with FcaRI (Gunn and Alter 2016). NK degranulation was maintained in the pregnant group despite their antibodies exhibiting lower binding affinity for FcγRIIIa (the cellular receptor that mediates NK degranulation). These observations further support the idea of compensatory mechanisms of the humoral immune system to offset pregnancy-intrinsic changes and maintain a healthy state.

Vastly different glycosylation of total and influenza-specific antibodies was observed in pregnant women compared to non-pregnant controls. The total Fc glycan profile in pregnant women, which included fucosylation and sialylation, was oriented towards reduced inflammation. In contrast, total Fab glycans, which involved increased fucosylation, agalactosylation, and bisection, were associated with increased inflammation. Non-inflammatory Fc galactosylation and sialylation as well as Fab N-acetylglucosamine bisection of bulk antibodies during pregnancy have been reported elsewhere (Bondt et al. 2014; Jansen et al. 2016; Bondt et al. 2016; Ruhaak et al. 2014; van de Geijn et al. 2009; Selman et al. 2012). Surprisingly, the Fc and Fab glycan profile of HA-specific antibodies were largely concordant within each group and included digalactosylated (G2) structures with increased sialylation and reduced bisection in the pregnant group, whereas agalactosylated, monosialylated, and bisected structures were predominant in the non-pregnant group. The association patterns remained broadly unchanged in the two groups, which indicates a coordinated global regulation of Fc and Fab glycosylation. However, other factors governing production of Fab and Fc glycans may reduce the overall level of coordination to reflect the putative greater level of Fab glycosylation during pregnancy (van de Bovenkamp et al. 2016).

Two distinct populations emerged in the PLSDA analysis, with Fc and Fab glycosylation being primary discriminatory elements; the clear separation of the two groups confirms the robustness of the data. The similarity in Fc and Fab glycosylation for the influenza-specific antibodies suggests some level of coordination of post-translational modifications in the Golgi that differs according to the B cell population. This is the first demonstration of Fab and Fc glycosylation of vaccine-specific antibodies consistent with non-inflammatory activity in pregnant women and different from the more inflammatory pattern in the non-pregnant group. Despite the disparity in antibody glycosylation, all the relevant Fc-mediated influenza-specific cellular functions were detected in the pregnant group.

We have previously shown that digalactosylated antibodies involved in NK-degranulation were enriched in cord blood; these digalactosylated antibodies preferentially bind to the neonatal Fc receptor (FcRn), which transits antibodies across the placenta (Jennewein et al. 2019; Martinez et al. 2019). Digalactosylated antibodies are reportedly more effective at facilitating NK-degranulation, a function the neonatal immune system is better equipped to deploy as opposed to phagocytosis or other non-neutralizing functions (Jennewein et al. 2019). For the mother, digalactosylation of antibodies during gestation may reflect another compensatory mechanism that maintains ADNP unaltered when other cellular functions may be reduced. Likewise, sialylation has been exploited in immunoglobulin therapy to endow anti-inflammatory benefits (Anthony et al. 2008; Pagan, Kitaoka, and Anthony 2018; Bruckner et al. 2017). Here, enhanced Fc-sialylation during pregnancy may contribute to dampened antibody-mediated inflammation consistent with the tolerizing state of pregnancy (Pagan, Kitaoka, and Anthony 2018).

Lastly, serum antibodies from pregnant women were retained longer in mouse circulation as demonstrated in *in vivo* passive transfer experiments, suggesting that beyond impacting the inflammatory state, the biophysical changes described above, in particular concomitant increased galactosylation and sialylation (Bas et al. 2019) may be intended to ensure sufficient circulating antibody during gestation during extended periods for protection of both the mothers and newborns. Together, the shifts observed in influenza-specific antibodies during pregnancy are consistent with reduced inflammation, more efficient Fc receptor engagement to maintain anti-viral function (as the magnitude of antibody available might be reduced), enhancement of placental transfer, reduction in antibody catabolism, and enrichment of antibodies that are more efficient for the newborn. The origin, regulation, and evolution of these pregnancy-associated antibody changes remains to be investigated.

Rather than broad immune suppression during pregnancy (Memoli et al. 2013), our data suggest that humoral immunity is selectively modulated by changes in antibody levels, avidity, FcR binding capacity, glycosylation patterns, and innate cellular functions. Our results also argue against universally compromised humoral immunity and in favor of discrete, antigen-dependent modifications.

While our study is limited by a small sample size and the retrospective analysis of immune responses to influenza seasonal vaccine strains, it still provides a valuable snapshot of vaccine-specific immunity in the pregnant population. Maternal infections place women and their infants at high risk. Understanding pregnancy-associated changes in antibody production, post-translational modifications and function, as well as how these processes affect vaccine responses will benefit vaccine design and will inform the development of much needed preventive and therapeutic strategies, particularly those for use in emergency situations, to improve maternal and infant health.

## Acknowledgments

H. X. was supported by the fund of FDA Office of Women’s Health.

## Author contributions

Conceptualization, M.F.J., H.X., G.A., and M.F.P.; Clinical samples: M.F.P, W.H.C and J.D. C. Investigation; M.F.J., M.K., F.J.N., P.R., and C.M.B.; Resources, H.X., G.A., and M.F.P.; Writing-Original draft, M.F.J, H.X., G.A., and M.F.P.; Writing-review and editing, J.D.C, W.H.C.; Supervision, H.X., G.A., and M.F.P.

## Figure Legends

**Supplemental Table 1:** Demographics of study participants.

|  | Non-pregnant | Pregnant |
| --- | --- | --- |
| **Age** | **31.35**<br>(Range: 23-44) | **27.64**<br>(Range: 18-35) |
| **Hormonal birth control** | **10:10**<br>(yes: no) | **N/A** |
| **Received flu vaccine** | **15:5**<br>(yes:no record) | **11:3**<br>(yes:no record) |
| **Received TDAP vaccine** | **0:20**<br>(yes:no) | **14:0**<br>(yes:no) |
| **Months between vaccine and sample collection** | **4.2**<br>(Range: 0-6) | **4.3**<br>(Range:2-6) |

## METHODS

### Human samples

Serum samples were obtained from pregnant (end of 3^rd^ trimester) and non-pregnant women who received egg-based inactivated influenza vaccine during the 2017-2018 flu season; samples were obtained within 4 months of vaccination **(Supplemental table 1, Supplemental figure 1)**. The vaccine components for the northern hemisphere 2017-2018 season influenza vaccine included H1N1 A/Michigan/45/2015, H3N2 A/Hong Kong/4801/2014, B/Phuket/3073/2013, and B/Brisbane/60/2008-like viruses((WHO) 2017). All subjects were HIV, HBV, and HCV negative and provided written informed consent before enrollment. This study was approved by the University of Maryland, Baltimore Institutional Review Board (HP-00065842 and HP-00040025) and by Partners Human Research Committee (Approval Number 2018P001060).

### Cell lines

Madin-Darby canine kidney-SIAT1 cells (SIAT1-MDCK, Sigma Aldrich, St. Louis, MO, USA), were grown in MEM plus 50 µg/ml of G418, 5% fetal bovine serum, 1X GlutaMAX™ and penicillin/streptomycin. THP-1 cells (ATCC, Manassas, VA, USA), were grown in R10 (RPMI plus 10% FBS L-glutamine and penicillin/streptomycin) supplemented with 0.01% β-mercaptoethanol.

### Viruses

H1N1 A/California/07/2009 (H1 CA), H1N1 A/Michigan/45/2015 (H1 MI), and H3N2 A/Hong Kong/4801/2014 (H3 HK) were grown in 9-10 day old embryonated eggs. Infectious viral titer, i.e., 50% tissue culture infectious dose (TCID50), was determined using a nucleoprotein-based enzyme-linked immunosorbent assay (ELISA) (Kosikova et al. 2018).

### Recombinant proteins

For titer and avidity determination, genes coding H3 HK HA, chimeric HA bearing H1 A/Puerto Rico/8/1934 stalk (group 1) with mismatched H16 HA head and chimeric HA bearing H3 HK stalk (group 2) with mismatched H4 HA head were synthesized as previously described (Stevens et al. 2004; Pica et al. 2012; Radvak et al. 2021) and were subcloned into the pFastBac gp67 vector (GenScript, Nanjing, China). Recombinant H3 HK HA, chimeric group 1 or group 2 stalk proteins expressed in Sf9 cells were column purified (GenScript). For functional assays and luminex Fc receptor binding and titer, recombinant H1 MI (HAΔTM H1N1 A/Michigan/45/2015), H1 CA (HAΔTM A/California/07/2009) and H3 HK proteins (H3ΔTM H3N2 A/Hong Kong/4801/2014) expressed in 293 cells were purchased from Immune Technology Corp. (New York, NY, USA).

### Anti-HA ELISA and avidity

HA-specific ELISA and avidity assays were performed as described (Kosikova et al. 2018). Briefly, serum samples were serially diluted and added to MaxiSorp microtiter plates (ThermoFisher) coated with 0.2 μg/mL of recombinant protein. Plates were incubated at room temperature for 60 min. For avidity determination, antigen-antibody complexes were dissociated by incubation with 100 μl of 4 M urea for 15 minutes and plates blocked again for an additional hour (Kosikova et al. 2018). Bound antibodies were detected by addition of horseradish peroxidase-labeled goat-anti-human IgG (Invitrogen, Carlsbad, CA, USA) followed by 1-Step Ultra TMB (ThermoFisher) substrate. Absorbance at 450 nm was measured in a Victor V multilabel reader (PerkinElmer, Waltham, MA, USA). Antibody titers were determined by interpolation in a Sigmoidal four parameter logistic regression curve. Avidity index was calculated based on the area of the entire antibody titration curve as described previously (Kosikova et al. 2018).

### Hemagglutination inhibition (HAI)

Sera were pre-treated with receptor-destroying enzyme (RDE) (Denka-Seiken, Tokyo, Japan), serially diluted two-fold (starting at 1:10) and incubated with 4 HA units per 25 μl of viruses at room temperature for 30 min. Hemagglutination was determined using 0.5% turkey red blood cells for H1N1 viruses or 0.75% guinea pig red blood cells in the presence of 20 nM oseltamivir for H3N2 viruses as described previously (Xie et al. 2015; Xie et al. 2011). HAI titers were expressed as the reciprocal of the highest serum dilution that resulted in complete HAI. A titer of 5 was assigned if no inhibition was observed at the starting dilution.

### Microneutralization (MN)

H1N1 and H3N2-specific MN titers were determined using a nucleoprotein-based ELISA as previously described (Kosikova et al. 2018). Briefly, RDE-treated sera were incubated with 100 50% tissue culture infectious dose (TCID_50_) of virus for 1 h at room temperature, 5% C0_2_, and then overlaid onto SIAT1-MDCK cells (Sigma) overnight. Infected cells were detected using influenza A nucleoprotein-specific monoclonal antibodies (Millipore, Burlington, MA, USA). MN titers were reported as the reciprocal of the highest serum dilution with ≥50% neutralization.

### Phagocytosis

The antibody-dependent monocyte phagocytosis assay (ADCP) used was adapted from Ackerman et al. 2011 and the antibody-dependent neutrophil phagocytosis assay (ADNP) was performed as described (Karsten et al. 2019). Briefly, influenza antigens, recombinant HAΔTM H1N1 A/Michigan/45/2015 and HAΔTM H3N2 A/Hong Kong/4801/2014 were biotinylated with a 20-fold excess of 10 mM EZ-Link NHS-LC-biotin (ThermoFisher) and unbound biotin was removed using a Zeba Spin Desalting Column (ThermoFisher). Antigens were then mixed with yellow-green fluorescent NeutrAvidin microspheres (ThermoFisher) at a ratio of 10 μl of beads per 10 μg of biotinylated protein and incubated at 37°C, 5% CO_2_ for 2 hours. Following incubation, beads were washed twice with PBS-0.01% BSA and resuspended in 1 mL of PBS-0.01% BSA. Serum samples were diluted 1:10, 1:50, and 1:100 in PBS, and 10 μl of each dilution was mixed with 10 μl of antigen-coupled fluorescent beads in a 96-well round bottom plate and incubated at 37°C, 5% CO_2_, for 2 h. The beads were then washed with 200 μl of PBS. For the monocyte phagocytosis, THP-1 cells (2.5×10^4^ cells in 200 μl per well) were added to wells containing the beads, and plates were incubated for 16 h at 37°C, 5% CO_2_. For the neutrophil phagocytosis, fresh neutrophils were isolated from whole blood of healthy adults and added (5×10^5^ cells in 200 μl) to bead-containing wells. Plates were incubated for 1h at 37°C, 5% CO_2_. Following incubation, cells were spun down and stained with anti-CD66b (clone:G10F5, BioLegend, Dedham, MA, USA) for 15 min. For both assays, cells were fixed with 100 μl of 4% paraformaldehyde (PFA) for 10 min, resuspended in 50 μl of PBS and analyzed by Flow cytometry using an iQue Screener PLUS (IntelliCyt, Albuquerque, NM, USA) and ForeCyt software (IntelliCyt). Influenza-specific IgG and PBS were included as positive and negative controls. Phagocytosis scores were calculated as the percentage of bead positive cells, multiplied by the geometric mean fluorescent intensity of the beads for bead positive cells, divided by 1×10^7^. All reported values are the area under the curve of the mean of two replicates.

### Complement deposition

Antibody-dependent complement deposition (ADCD) was performed as previously described (Fischinger et al. 2019). Biotinylated recombinant HAΔTM H1N1 A/Michigan/45/2015 and H3ΔTM H3N2 A/Hong Kong/4801/2014 were coupled to red fluorescent NeutrAvidin microspheres (ThermoFisher), incubated with sera and added to single wells as described above. Lyophilized guinea pig complement (Cedarlane, Burlington, ON, Canada) was resuspended in ice-cold water and diluted 1:100 in cold veronal buffer supplemented with 0.1% fish skin gelatin (Boston Bio Products, Boston, MA, USA). 200 μl of diluted complement was added to each well, and plates were incubated for 20 min at 37°C, 5% CO_2_. Plates were then washed twice with ice-cold 15 mM EDTA-PBS and stained with anti-guinea pig C3-FITC (MP Biomedical, Santa Ana, CA, USA). After a final wash, beads were re-suspended in 50 μl of PBS and analyzed on the iQue Screener PLUS. Red-fluorescent beads were gated based on size. Complement deposition was determined as the geometric mean fluorescent intensity of red fluorescence of the beads divided by 1,000. Data presented is area under the curve of the complement score averaged for two replicates.

### NK cell Activation

ELISA plates were coated with recombinant HAΔTM H1N1 A/Michigan/45/2015 and H3ΔTM H3N2 A/Hong Kong/4801/2014 at 3 μg/mL of PBS for 2 h at 37°C, 5% CO_2_. Plates were then washed three times with PBS and blocked overnight with 200 μl of 5% BSA in PBS. The next day, plates were washed, and samples were added in three serial dilutions (1:10, 1:50, 1:100) in a 50 μl volume and incubated 2 h at 37°C, 5% CO_2_. NK cells were isolated from healthy adult PBMC using a RosetteSep NK Cell Enrichment Kit (Stem Cell Technologies, Vancouver, BC, Canada), according to manufacturer’s instructions followed by density gradient centrifugation in Histopaque. A suspension of 1.5×10^6^ cells/ml in R10 was prepared, supplemented with 1 ng/ml rhIL15 and incubated overnight at 37°C, 5% CO_2_. Prior to the assay, a suspension of 2.5×10^5^ cells/ml was prepared and 2.5 μg/ml of Brefeldin A (Biolegend) and Golgistop (BD Biosciences, Franklin Lake, NJ, USA) and 2.5 µl of anti-CD107a (Clone:555802, BD Biosciences) were added. NK cells (5×10^4^ in 200μl) were added to each well containing Ag-Ab, and plates were incubated for 5 h at 37°C, 5% CO_2_. Following the incubation and washing, cells were stained with anti-CD3 (Clone:UCHT, BD Biosciences), anti-CD56 (Clone: B159, BD biosciences), and anti-CD16 (Clone: 3G8, BD biosciences), washed, fixed, permeabilized with FIX&Perm kit (ThermoFisher), and stained intracellularly with anti-IFNγ (Clone:B27, BD biosciences) and MIP-1β (Clone:D21-1351, BD Biosciences). Cells were resuspended in PBS and analyzed on the iQue Screener as described above. Influenza-specific IgG and PBS (diluent) were included as positive and negative controls. Negative control wells were used to set gates. Data was reported as the percentage of CD16+CD56+CD3-NK cells positive for CD107a, IFNγ, or MIP-1β and is the area under the curve of the mean of three replicates divided by 100,000.

### Cellular cytotoxicity

Antibody-dependent cellular cytotoxicity (ADCC) assay was performed as previously described (Jacobsen et al. 2017) with modifications. Briefly, SIAT1-MDCK cells were seeded onto 96-well white flat-bottom plates (PerkinElmer), infected with H1 MI or H3 HK viruses and incubated for 6 h at 37°C. RDE-treated sera (25 µl/well) and Jurkat effector cells (Promega, Madison, WI, USA) (25 µl/well) were added to infected cells and incubated overnight at 37°C. Bio-Glo luciferase assay reagent (Promega) was then added, and luminescence was measured using a Victor V multilabel reader (PerkinElmer).

### Antibody subclass and FcR binding

IgG, IgA, IgM, and Fc receptor binding analyses were performed using a multiplex luminex assay as previously described (Brown et al. 2012). Briefly, recombinant HAΔTM H1N1 A/Michigan/45/2015 and H3ΔTM H3N2 A/Hong Kong/4801/2014 were coupled to MagPlex microspheres (Luminex corporation, Austin, TX, USA) at a ratio of 25 μg of protein to 400 μl of beads. Microspheres were activated with 100 mM monobasic sodium phosphate pH 6.2 in the presence of 50 mg/mL EDC and 50 mg/ml sulfo-NHS, washed with 0.05 M 2[N-Morpholino]ethanesulfonic acid (MES) pH 5.0, incubated with antigen for 2 h at 37°C, 5% CO_2_, and blocked with PBS-TBN (0.1% BSA, 0.02% tween-20, 0.05% Azide in PBS, pH 7.4) for 30 min. The beads were washed in PBS-0.05% tween-20, resuspended in 250 μl of PBS. Beads were added to 384-well plates and incubated with sera at a final 1:500 dilution in luminex wash buffer (PBS-0.05% BSA-0.001% tween-20) for 2 h at room temperature and with shaking (800 rpm).

To detect antibody subclasses, following incubation, beads were washed in luminex buffer and incubated with PE-labeled anti-IgG, -IgG1, -IgG2, - IgG3, -IgG4, -IgA1, -IgA2, and -IgM (Southern Biotech, Birmingham, AL, USA) for 1 h at room temperature with shaking. After washing, beads were resuspended in Qsol buffer (IntelliCyte) and plates were read on an iQue Screener PLUS. H1N1 and H3N2 coupled beads were distinguished based on size and granularity, and data were analyzed as median fluorescent intensity of PE for each bead population.

To investigate Fc receptor binding recombinant Fc receptors with an AviTag were biotinylated using a Bir500 kit (Avidity, Aurora, CO, USA) according to manufacturer’s instructions and purified using a Zeba Spin Desalting Column. Biotin-labeled FcRs were then incubated with streptavidin-PE (Prozyme, Hayward, CA, USA) for 10 min and excess streptavidin was quenched with 20 μM biotin (Avidity) for another 10 min. Following bead and serum incubation, PE-Fc receptors were then added and incubated for a further 1 h at room temperature with shaking. Plates were then washed and analyzed on an iQue Screener as described above. Data reported for each sample represents the mean of two replicates.

### Glycosylation analysis

Antigen-specific glycosylation analysis was performed as described in Jennewein et al. 2019. Briefly, recombinant HAΔTM H1N1 A/Michigan/45/2015 and H3ΔTM H3N2 A/Hong Kong/4801/2014 were biotinylated and coupled to streptavidin magnetic beads (New England Biolabs) at a ratio of 2.5 μg of protein to 25 μl of beads. Sera (200 μl) were first incubated with non-coated NeutrAvidin beads to remove non-specific binding and then added to the antigen-coated beads and incubated for 1 h at 37°C. Antibodies were eluted by incubation in 50 μl of pH 2.9 citrate buffer for 30 min at 37C. Samples were then spun down and the eluted antibodies, contained in the supernatant, were neutralized with 30 μl pH 8.9 potassium phosphate buffer. Antibodies were then coupled to protein G beads (New England Biolabs). After beads were washed, IDEZ was used to cleave the Fab (in the supernatant) from the Fc (remained on the magnetic beads) (New England Biolabs) for 1 h at 37°C and collected. The two fragments were deglycosylated and fluorescently labeled using a GlycanAssure APTS kit (ThermoFisher scientific) according to manufacturer’s instructions. For the Fc fragment, which remained bound to the protein G beads, an additional magnetic separation after PNGase glycan cleavage separated the glycans from remaining magnetic beads and the protocol then proceeded per manufacturer’s instructions. Glycans were analyzed on a 3500xL genetic analyzer. Glycan fucosyl and afucosyl libraries (Prozyme) were used to assign 24 discrete glycan peaks using GlycanAssure software (ThermoFisher). Data are reported as percentages of total glycans for each of the glycan peaks.

### In vivo Ab retention

Pooled sera from pregnant and non-pregnant women were transferred intraperitoneally into naïve female Balb/c mice (Charles River, Wilmington, MA) at 0.2 ml/mouse. The circulating concentration of human IgG after passive transfer was measured by a Human IgG Easy Quantification ELISA Kit (Cell Sciences, Newburyport, MA). A human IgG calibrated standard was included for calculation of IgG content.

### Statistical Analysis

Mann-Whitney test (univariate analysis) was used to examine differences in readouts between pregnant and non-pregnant groups. P values are two-sided, *p<0.05, **p<0.01, ***p<0.001, and ****p<0.0001. Data on radar plots were z-score normalized individually along each radar to an average of 0 with a standard deviation of 1. Multivariate Principal Component Analysis (PCA) and Least absolute shrinkage and selection operator LASSO Partial Least Squares Discriminant Analysis (LASSO-PLSDA) were used to distinguish pregnant vs. non-pregnant immune profiles. LASSO is a regression analysis that selects variables that contribute to the differences between samples arranged along an ordinal dependent variable. It selects features that are required to create a maximum separation which are then visualized on the PLSDA plot. Missing values were imputed using the k-nearest neighbor. Variables were centered and scaled to a standard deviation of 1. PCA was used to visualize if pregnant and non-pregnant samples separate into discrete groups within the two-dimensional principal component (PC) space. The LASSO-PLSDA model had a calibration success of 100% with a leave-1-out CV success of 97.0588%. Univariate analysis was performed with Prism 7 (GraphPad, San Diego, CA). MATLAB was used to perform all multivariate analysis.

---

**Supplemental Figure 1:**
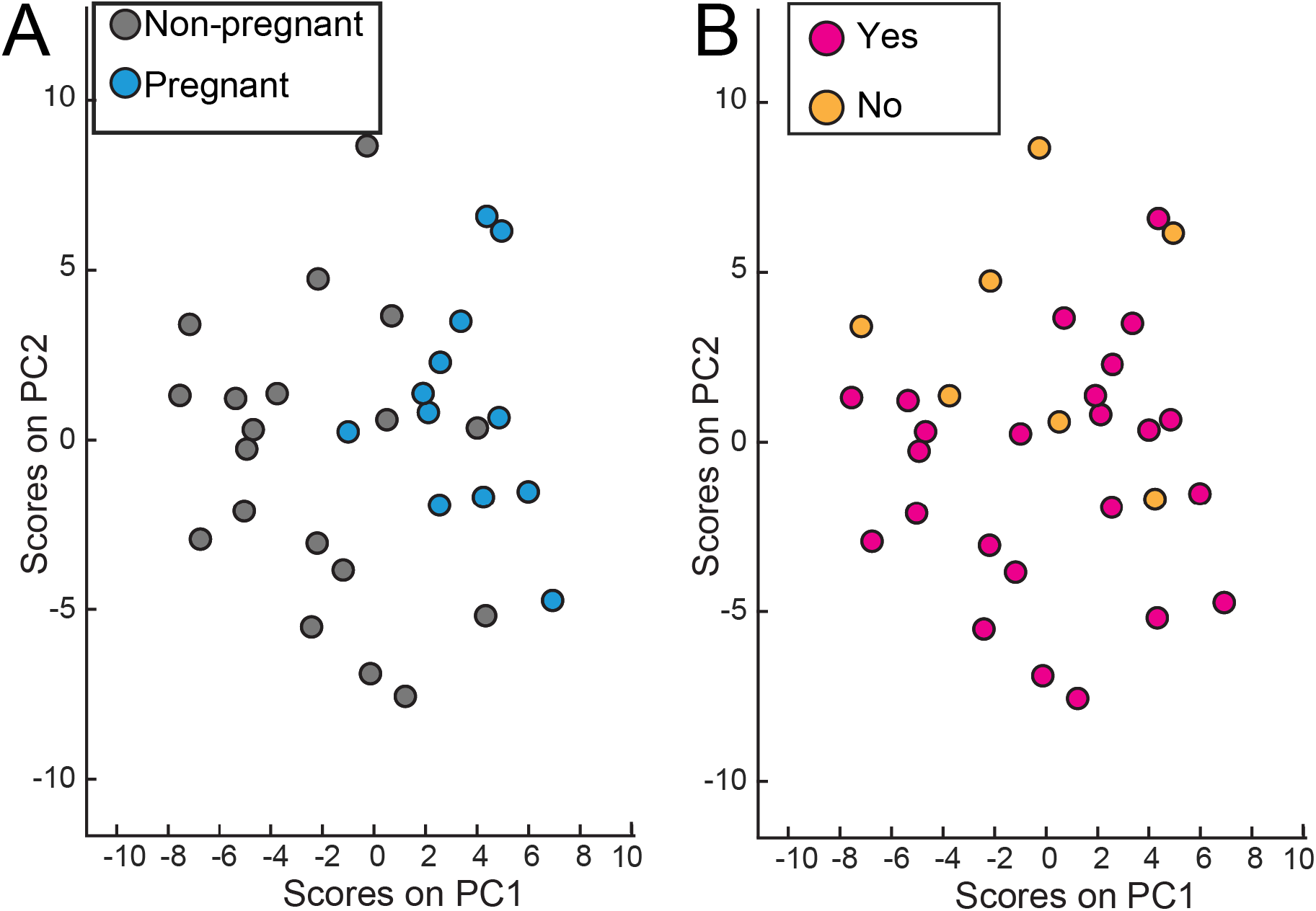
Relationship of antibody profiles and vaccination. PCA analysis was used to distinguish effects of vaccination. No differences were observed across the antibody profiles for vaccination and un-vaccinated participants. Principle component 1 (PC1) contains 12% of the variance and PC2 contains 10.5% of the variance. **A**. Dots colored to distinguish pregnant (blue dots) and non-pregnant (grey dots) women. **B**. Dots colored to distinguish women who received seasonal influenza vaccine (pink dots) and women without clinical record of vaccination (orange dots).

